# Anodal transcranial direct-current stimulation over the left dorsolateral prefrontal cortex and perception of emotional faces

**DOI:** 10.1101/2020.02.09.924910

**Authors:** Joaquim Brasil-Neto, Tamiris Bernardes, Camila Rosa da Silva, Aline Iannone, Ana Garcia, Maria Clotilde Henriques Tavares

**Affiliations:** Department of Physiological Sciences, Biology Institute, University of Brasília, Darcy Ribeiro Campus, Brasília, DF, Brazil

**Keywords:** tDCS, emotional stimuli, response times

## Abstract

The prefrontal cortices have been shown to be engaged in the perception of emotional faces, and imaging studies correlate activation of the left dorsolateral prefrontal cortex (l-DLPFC) with correct identification of emotional expressions. Transcranial direct current stimulation (tDCS)has been shown to modulate cortical function and might influence the perception of emotional faces. Our aim was to evaluate the possible effects of tDCS on a task that required correct identification of emotions represented by human faces presented on a screen. Ten volunteers (ages 18-30; mean = 23.2; SD = 4.3) were evaluated. The experiments were carried out in two 20 minutes stimulation sessions (real and sham tDCS) 4 days apart, at least, to avoid carryover effects, and session orders were randomized. The anode was placed over the l-DLPFC (F3 of the international 10-20 EEG electrode positioning system) and the cathode over the contralateral supraorbital region. Emotional stimuli were 120 consecutive human faces to be verbally classified as “happy”, “surprised”, “neutral”, “angry” or “sad”. Real anodal tDCS over the l-DLPFC resulted in significantly longer response times with a trend towards increased error rates, compared to the sham condition.

## Introduction

Impaired detection of emotional social cues is present in certain neuropsychiatric conditions. Although the prefrontal cortices have been shown to be engaged in facial emotion recognition, the exact roles of the left and right dorsolateral prefrontal cortices is controversial.

Transcranial Direct Current Stimulation (tDCS) is a noninvasive brain stimulation technique that allows temporary modulation of neuronal activity in specific cortical regions and can influence subject performance in behavioral tasks (1). It has also been proposed as a neuromodulatory tool with potential therapeutic uses, since tDCS sessions might modulate neural funtion beyond stimulation sessions (2). Therefore, tDCS neuromodulation sessions might be useful in the treatment of conditions in which there is impairment in processing of emotional faces. However, the neural targets and whether they should receive inhibitory or excitatory stimulation are still matters of speculation.

Several studies have shown that tDCS excitatory (anodal) activation over the left dorsolateral prefrontal cortex (l-DLPFC) can improve purely cognitive (“cold”) executive functions (EF) in healthy subjects and patients (3–7), but its effect upon affective-related (“hot”) EF is still unclear (8).

Many studies have shown that the DLPFC is activated during processing of emotional information.

In a study using functional magnetic resonance imaging (fMRI), Sergerie et al (9) showed that l-DLPFC activation was associated with successful encoding of faces with an emotional expression, and activity in the right DLPFC with memory for faces, regardless of emotional expression. However, a tDCS study involving simultaneous stimulation of both prefrontal cortices (10), found that anodal right/cathodal left stimulation (therefore probably resulting in l-DLPFC inhibition and r-DLPFC stimulation) was able to speed up responses to threatening faces, although only in male subjects. Stimulation of the l-DLPFC by means of anodal tDCS has been shown to facilitate attentional disengagement away from emotional faces (happy, disgusted and sad expressions), whereas active stimulation of the r-DLPFC slowed gaze disengagement to the same emotional stimuli (11).

Yang et al.(12) have recently reported that anodal tDCS over the r-DLPFC improved correct detection of schematic emotional faces, but only during the tDCS session, with no effect when tests were done immediately after 15 min of tDCS at 1.5 mA. Moreover, the improvement was only seen for faces expressing positive emotions. This is in contrast to the current view that the r-DLPFC would be superior in recognition of all emotions, or superior in the recognition of negative emotions (13).

As already mentioned, l-DLPFC activation in neuroimaging studies has been shown to be correlated with correct encoding of facial expressions. Therefore, we have hypothesized that l-DLPFC anodal stimulation might be capable of facilitating correct classification of emotional faces (“happy”, “surprised”, “neutral”, “angry” or “sad”) in young normal volunteers. Moreover, such effect should be persistent after individual tDCS sessions, thus indicating a potential for therapeutic use. In this study, we set out to test this hypothesis.

## Methods

Ten healthy subjects, 4 men and 6 women, with a mean age of 23.2 years (range 18-30 years) participated in the study, which was approved by the local ethics committee. All were right-handed with normal or corrected-to-normal vision. All participants were blind to the purpose of the experiments. Before being enrolled in the study, all volunteers were screened for depression and anxiety (Beck scales) and refrained from smoking as well as from ingestion of any caffeinated beverages. All denied use of psychoactive drugs or medications. Each subject participated in two experimental sessions, which were done at least four days apart, in order to avoid carry-over effects. Each experimental session involved 20 minutes of either real or sham tDCS for 20 minutes, during which subjects sat comfortably and watched a short movie, followed by the emotional faces recognition task. A crossover design was employed, and half the subjects received sham tDCS on the first session whereas the other half received real stimulation.

### Face recognition task

Visual stimuli were presented on a 17-inch laptop screen.They consisted of human faces extracted from Minear and Park’s database (14). The screen resolution was 1024 X 768, and the refresh rate was 100 Hz. Participants responded using a microphone and the bar space and responses were recorded and timed by a software, with subsequent manual review. During each session, a total of 120 emotional faces were presented, with the following emotions: "“happy”, “surprised”, “neutral”, “angry” and “sad”. Presentation times and screen locations were randomly changed, in order to avoid a learning effect. One third of the stimuli had positive valences, one third negative valences, and one third were neutral. Volunteers were instructed to verbally name the perceived emotion, as fast and accurately as possible, while simultaneously pressing the space bar. Both the number of correct responses and the response times were recorded for each individual.

### Transcranial direct-current stimulation

During the experimental sessions, subjects were comfortably seated and were asked to remain still and to refrain from talking, while watching a short movie. In each subject, electrodes (35 cm2 saline-soaked sponges covering conductive rubber pads) were firmly attached to the skull with the aid of Velcro straps. The skin was not abraded, and no other conductive media besides saline solution were used. The anode was placed over the F3 position of the 10-20 international system and the cathode (“reference”) over the right supraorbital region. A commercially available continuous current stimulator (TransCranial®, Hong Kong) was used to apply a 2mA current for 20 minutes. In the sham condition, the current was initially turned on for 10 seconds and then gradually turned off over the course of 10 seconds, so that the tingling sensation usually felt at the beginning of real stimulation would be perceived by the participant.

## Results

All tDCS sessions were well tolerated by all subjects, without discomfort or unwanted side effects.

Response times did not have a normal distribution, as evidenced by the Shapiro-Wilk test; therefore, non-parametric statistics was used to analyze the data (Mann-Whitney U-Test and Kruskal-Wallis Test).

When comparing all responses, there was a significant increase in response times after real tDCS: mean response times were 716.2 ms (SEM 6.54) for real tDCS and 705.5 ms (SEM 6.29) for sham (p< 0.05) (Figure 1). There was also a non-significant trend towards more errors in the real condition: mean emotion classification error rates were 11.75 % for real and 10.84 % for sham (p=0.61). There were no differential effects on specific emotions (Figures 2, 3).

**Fig. 1.**
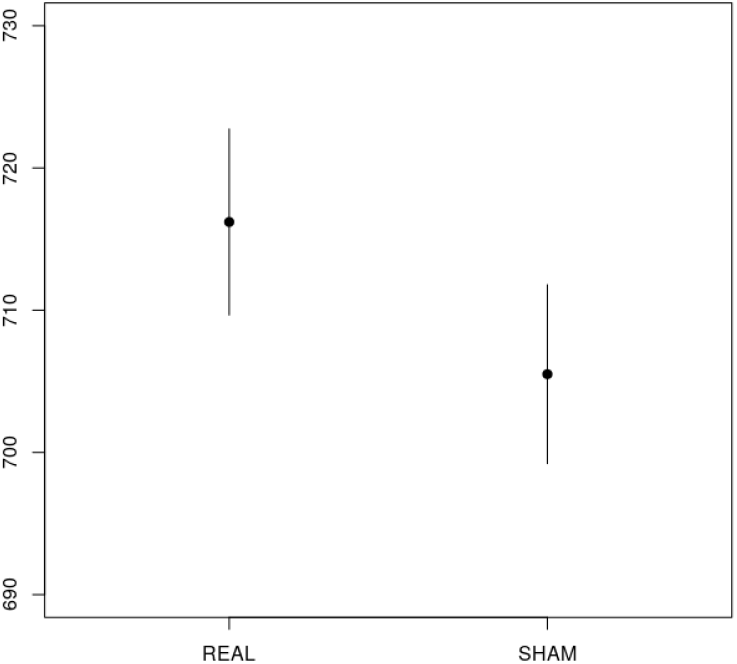
Response times (msec) were significantly longer after real tDCS compared to the sham condition. Error bars are standard deviations of the mean.

**Fig. 2.**
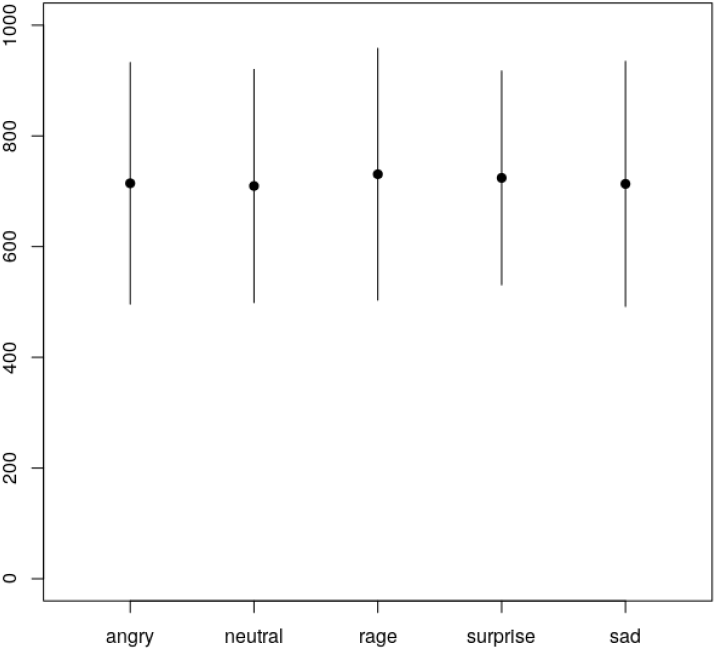
Mean response times for each emotion under real stimulation.Error bars are standard deviations of the mean.

**Fig. 3.**
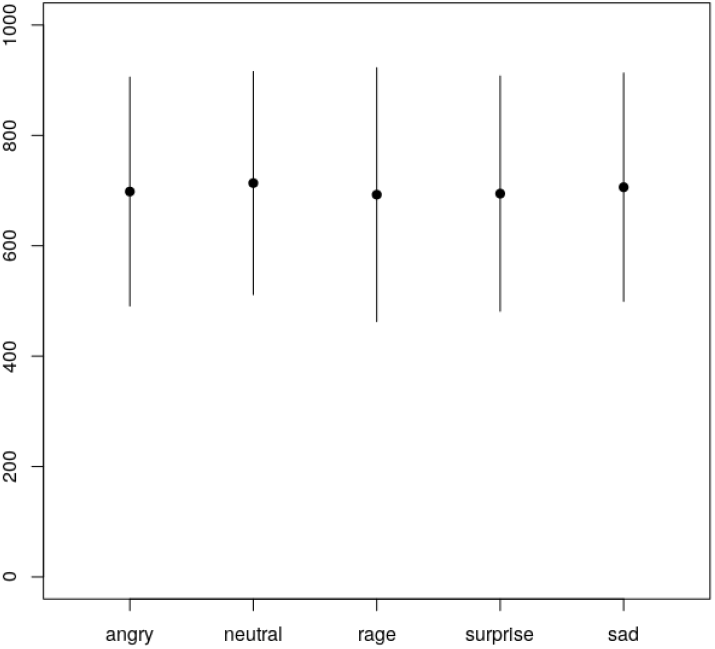
Mean response times for each emotion under sham stimulation.Error bars are standard deviations of the mean.

## Conclusions

Contrary to our experimental hypothesis, anodal tDCS over the l-DLPFC resulted in a certain degree of impairment in detection of emotional faces, with a significant increase in response times and a trend towards more classification errors. These results, however, are in line with neuroimaging studies (9), which have demonstrated activation of this area during encoding of faces with emotional expressions. The fact that in this study anodal tDCS has apparently disrupted l-DLPFC function is not totally unexpected, since other authors have already pointed out that the usual dichotomy anodal-excitatory and cathodal-inhibitory is not invariably true (15–17).

Since each subject was exposed to both conditions (sham and real tDCS) in a cross-over design, confounding factors due to inter-individual variability were minimized.

This study involved a small number of subjects, but the results obtained so far point to a role for the l-DLPFC in processing of emotional faces. However, no differential effects were seen for the emotions under study. The study of a larger number of subjects might shed light on the significance of the observed increase in emotion identification errors.

The position of the reference electrode on the contralateral supraorbital area has been a common one, even in neurocognitive studies (18). However, to rule out unintended stimulation of the r-DLPFC it would be interesting to use a cathode with an area of 100 cm2, as this has been shown to induce much weaker currents, rendering the cathode physiologically inactive.

Finally, performance changes occurred off-line, i.e., after a tDCS session. This is important, since any clinical use of transcranial electrical stimulation for modulation of neuronal function has to rely on its after-effects between sessions.

## ACKNOWLEDGEMENTS

This work was supported by a grant from FAPDFFundação de Apoio à Pesquisa do Distrito Federal, registration number 0193.001401/2016.

## Bibliography

1. Paulo Sérgio Boggio, Camila Campanhã, Cláudia A Valasek, Shirley Fecteau, Alvaro Pascual-Leone, and Felipe Fregni. Modulation of decision-making in a gambling task in older adults with transcranial direct current stimulation. Eur. J. Neurosci., 31(3):593–597, 2010.

2. Jean-Pascal Lefaucheur, Andrea Antal, Samar S Ayache, David H Benninger, Jérôme Brunelin, Filippo Cogiamanian, Maria Cotelli, Dirk De Ridder, Roberta Ferrucci, Berthold Langguth, Paola Marangolo, Veit Mylius, Michael A Nitsche, Frank Padberg, Ulrich Palm, Emmanuel Poulet, Alberto Priori, Simone Rossi, Martin Schecklmann, Sven Vanneste, Ulf Ziemann, Luis Garcia-Larrea, and Walter Paulus. Evidence-based guidelines on the therapeutic use of transcranial direct current stimulation (tDCS). Clin. Neurophysiol., 128(1): 56–92, January 2017.

3. Tino Zaehle, Pascale Sandmann, Jeremy D Thorne, Lutz Jäncke, and Christoph S Herrmann. Transcranial direct current stimulation of the prefrontal cortex modulates working memory performance: combined behavioural and electrophysiological evidence. BMC Neurosci., 12(1):2, January 2011.

4. Nadia Bolognini, Felipe Fregni, Carlotta Casati, Elena Olgiati, and Giuseppe Vallar. Brain polarization of parietal cortex augments training-induced improvement of visual exploratory and attentional skills. Brain Res., 1349:76–89, 2010.

5. Vincent P Clark, Brian a Coffman, Andy R Mayer, Michael P Weisend, Terran D R Lane, Vince D Calhoun, Elaine M Raybourn, Christopher M Garcia, and Eric M Wassermann. TDCS guided using fMRI significantly accelerates learning to identify concealed objects. Neuroimage, 59(1):117–128, November 2012.

6. Valentina Fiori, Michela Coccia, Chiara V Marinelli, Veronica Vecchi, Silvia Bonifazi, M Gabriella Ceravolo, Leandro Provinciali, Francesco Tomaiuolo, and Paola Marangolo. Transcranial direct current stimulation improves word retrieval in healthy and nonfluent aphasic subjects. J. Cogn. Neurosci., 23(9):2309–23, September 2011.

7. Felipe Fregni, Paulo S Boggio, Michael Nitsche, Felix Bermpohl, Andrea Antal, Eva Feredoes, Marco A Marcolin, Sergio P Rigonatti, Maria T A Silva, Walter Paulus, and Alvaro Pascual-Leone. Anodal transcranial direct current stimulation of prefrontal cortex enhances working memory. Exp. Brain Res., 166(1):23–30, 2005.

8. Vahid Nejati, Mohammad Ali Salehinejad, and Michael A Nitsche. Interaction of the left dorsolateral prefrontal cortex (l-DLPFC) and right orbitofrontal cortex (OFC) in hot and cold executive functions: Evidence from transcranial direct current stimulation (tDCS). Neuroscience, 369:109–123, January 2018.

9. Karine Sergerie, Martin Lepage, and Jorge L Armony. A face to remember: emotional expression modulates prefrontal activity during memory formation. Neuroimage, 24(2):580–585, January 2005.

10. Massimiliano Conson, Domenico Errico, Elisabetta Mazzarella, Marianna Giordano, Dario Grossi, and Luigi Trojano. Transcranial electrical stimulation over dorsolateral prefrontal cortex modulates processing of social cognitive and affective information. PLoS One, 10(5): e0126448, May 2015.

11. Alvaro Sanchez-Lopez, Marie-Anne Vanderhasselt, Jens Allaert, Chris Baeken, and Rudi De Raedt. Neurocognitive mechanisms behind emotional attention: Inverse effects of anodal tDCS over the left and right DLPFC on gaze disengagement from emotional faces. Cogn. Affect. Behav. Neurosci., 18(3):485–494, June 2018.

12. Tao Yang and Michael J Banissy. Enhancing anger perception in older adults by stimulating inferior frontal cortex with high frequency transcranial random noise stimulation. Neuropsychologia, 102:163–169, July 2017.

13. Giulia Prete, Bruno Laeng, and Luca Tommasi. Transcranial random noise stimulation (tRNS) over prefrontal cortex does not influence the evaluation of facial emotions. Soc. Neurosci., 14(6):676–680, December 2019.

14. Meredith Minear and Denise C Park. A lifespan database of adult facial stimuli. Behav. Res. Methods Instrum. Comput., 36(4):630–633, November 2004.

15. Sabrina Brückner and Thomas Kammer. Both anodal and cathodal transcranial direct current stimulation improves semantic processing. Neuroscience, 343:269–275, February 2017.

16. Liron Jacobson, Meni Koslowsky, and Michal Lavidor. tDCS polarity effects in motor and cognitive domains: a meta-analytical review. Exp. Brain Res., 216(1):1–10, January 2012.

17. Carlo Miniussi, Justin A Harris, and Manuela Ruzzoli. Modelling non-invasive brain stimulation in cognitive neuroscience. Neurosci. Biobehav. Rev., 37(8):1702–1712, September 2013.

18. Paulo S Boggio, Felix Bermpohl, Adriana O Vergara, Ana L C R Muniz, Fernanda H Nahas, Priscila B Leme, Sergio P Rigonatti, and Felipe Fregni. Go-no-go task performance improvement after anodal transcranial DC stimulation of the left dorsolateral prefrontal cortex in major depression. J. Affect. Disord., 101(1-3):91–98, 2007.

